# Individual neurophysiological signatures of spontaneous rhythm processing

**DOI:** 10.1101/2022.03.14.484286

**Authors:** A. Criscuolo, M. Schwartze, M.J. Henry, C. Obermeier, S.A. Kotz

## Abstract

When sensory input conveys rhythmic regularity, we can form predictions about the timing of upcoming events. Although rhythm processing capacities differ considerably between individuals, these differences are often obscured by participant- and trial-level data averaging procedures in M/EEG research. Here, we systematically assessed the neurophysiological variability displayed by individuals listening to isochronous equitone sequences interspersed with unexpected deviant tones. We first focused on rhythm tracking and tested the anticipatory phase alignment of delta-band activity to expected tone onsets. These analyses confirmed that individuals encode temporal regularities and form temporal predictions, but highlight clear inter- and intra-participant variability. This observation may indicate individual and flexible tracking mechanisms, which show consistency at the single-trial level, but variability over trials. We then modelled single-trial time-locked neural responses in the beta-band to investigate individual tendencies to spontaneously employ binary grouping (“tic-toc effect”). This approach identified binary (strong-weak), ternary (strong-weak-weak), and mixed accentuation patterns, confirming the superimposition of a basic beat pattern. Furthermore, we characterized individual grouping preferences and tendencies to use binary, ternary, or combined patterns over trials. Importantly, the processing of standard and deviant tones was modulated by the employed pattern. The current approach supports individualized neurophysiological profiling as a sensitive strategy to identify dynamically evolving neural signatures of rhythm and beat processing. We further suggest that close examination of neurophysiological variability is critical to improve our understanding of the *individual* and *flexible* mechanisms underlying the capacities to rapidly evaluate and adapt to environmental rhythms.

**Significance statement:** For decades, music, speech and rhythm research investigated how humans process, predict, and adapt to environmental rhythms. By adopting a single-trial and -participant approach, we avert the common pooling of EEG data in favor of individual time-varying neural signatures of rhythm tracking and beat processing. The results highlight large inter- and intra-individual differences in rhythm tracking, arguing against the typically documented phase-specificity for entrainment. On top of that, we characterize individual variability in beat processing, by showing that binary, ternary and other accentuation patterns are used over time, and ultimately affect the processing of (un-)expected auditory events. The approach aids individual neural profiling and may therefore allow identifying altered neural activity and its consequences in natural listening contexts.

## 1. Introduction

Due to the inherently rhythmic nature of many environmental stimuli, neurocognitive functions such as attention (Lakatos et al., 2008) sensorimotor behavior (Merker et al., 2009), speech (Giraud & Poeppel, 2012), reading (Goswami, 2011), and music (Doelling & Poeppel, 2015) rely on basic timing capacities. To generate a temporally coherent representation of a rhythmic environment, we track stimulus periodicities, use smart grouping, and continuously segment and combine multiple inputs in time (Buzsáki, 2009; Schroeder & Lakatos, 2009; Thut et al., 2012a; Zoefel & VanRullen, 2016). According to the dynamic attending theory (Large & Jones, 1999) these timing processes reflect how internal rhythms synchronize with external rhythms. This and similar theoretical views (Fraisse, 1963; p. 18) suggest that oscillatory brain activity instantiates a realistic model for such “adaptation by anticipation”. Accordingly, temporally regular sensory input would make future events predictable and thereby facilitate sensory processing, perception, allocation of attention, and the overall effectiveness of behavior (Friston, 2005; Arnal, 2012; Schröger et al., 2015; Koelsch et al., 2019).

However, *continuous change is a fundamental characteristic of life* (Schwartze et al., 2011), and next to regular rhythms, we frequently experience irregular rhythms or sudden changes in the environment. To account for these dynamics, any realistic adaptation mechanism likely tolerates a certain degree of temporal irregularity or unpredictability while trying to achieve synchronization (Barnes & Jones, 2000). Endogenous oscillatory activity must hence not only be precise and stable over time, but also flexible enough to achieve adequate adaptive timing (e.g., by speeding-up or slowing-down accordingly). Indeed, oscillatory brain activity can actively track and process (quasi-)periodic and never strictly isochronous signals such as speech, which includes rhythmic variations at phoneme (25-35Hz) to syllable (4-8Hz) and word (1-3Hz) rates (Giraud & Poeppel, 2012). Furthermore, neural oscillations can rapidly adapt to changes in the sensory environment, likely through phase resetting (Barnes & Jones, 2000; Haegens & Zion Golumbic, 2018; Mormann et al., 2005; Obleser et al., 2012; Zoefel et al., 2018).

Particularly, delta- (δ; 1-4Hz) and beta- (β; 12-25Hz) frequency-band neural oscillations have been associated with rhythm processing, temporal prediction, and attention in humans (Arnal, 2012; Biau & Kotz, 2018; Colling et al., 2017; Fujioka et al., 2012, 2015; Morillon et al., 2016; Nozaradan et al., 2015, 2017a) and in non-human primates (Bartolo & Merchant, 2015; Merchant et al., 2015; Merchant & Bartolo, 2018; Patel & Iversen, 2014). However, prior behavioural studies reported large within-subject variability in individual beat preferences across trials (Baath, 2015; Poudrier, 2020). Thus, a critical question arises: does individual neurophysiological variability increase the explanatory power of how individuals process the beat and rhythm (see Kononowicz & van Rijn (2015) for the case of duration estimation; Grahn & McAuley, 2009; Waschke et al., 2021).

To address this question, we let participants passively listen to isochronous (1.5Hz) equitone sequences, comprising frequent standard and either one or two amplitude-attenuated deviant tones while their EEG was recorded. Neural signatures of rhythm tracking were assessed by quantifying the trial-level consistency of delta-band phase alignment towards expected tone onsets. We expected to observe a predictive phase-alignment of the high-excitability phase towards tone onsets.

Next, we focused on the known human disposition to group two or three adjacent tones when listening to isochronous equitone sequences (Brochard et al., 2003). This results in perceived binary (strong-weak (S-w)) or ternary (S-w-w) accentuation patterns, ultimately resembling *beat processing*. In other words, auditory sequences are subdivided in regular groups of adjacent events according to a regular superimposed beat structure. Importantly, even with physically identical stimuli, this beat can influence observable behaviour and underlying neural activity (Abecasis et al., 2005; Baath, 2015; Brochard et al., 2003; Fujioka et al., 2015; Poudrier, 2020; Schmidt-Kassow et al., 2011). Informed by previous evidence showing a pivotal role of beta-band activity in beat processing (Fujioka et al., 2012; Fujioka et al., 2015), we expected to observe binary and ternary accentuation patterns in the envelope of beta-band activity. Unlike in previous studies, we zoomed into individual differences to investigate whether: (i) individuals always accentuate, (ii) individuals accentuate in a similar way, and (iii) accentuation patterns modulate deviant processing (Brochard et al., 2003). To this end, we modelled single-participant and single-trial time-locked fluctuations in the beta-band, aiming to characterize inter- and within-participant predispositions to superimpose beat-like accentuation patterns and opt for binary, ternary, and combined beat processing over trials. We expected this analysis approach to deliver insights into the intra- and inter-individual neurophysiological variability associated with *individual and flexible* mechanisms employed to evaluate and adapt to (un)predictable environmental rhythms.

## 2. Materials & Methods

### 2.1. Participants

Twenty native German speakers participated in the study and signed written informed consent in accordance with the guidelines of the ethics committee of the University of Leipzig and the declaration of Helsinki. Participants (9 females; 21-29 years of age, mean 26.2 years) were right-handed (mean laterality coefficient 93.8, Oldfield, 1971), had normal or corrected-to-normal vision and no known hearing deficits. Participants received 8€/hr for taking part in the study. Beyond daily music listening, the participants had no defined musical expertise.

### 2.2. Experimental design and procedure

The stimuli comprised 192 sequences, consisting of 13-to-16 400Hz, 50ms, 70dB SPL tones (standard; STD), presented in two recording sessions. One or two deviant tones (DEV), attenuated by 4dB relative to the STD tones, were embedded in each sequence replacing STD tones. The first DEV tone could either occur in an odd or even-numbered position (8-11^th^), while the second always fell on the 12^th^ position (Fig.1; corresponding to a hypothetical binary Strong-weak pattern; S-w). The inter-onset-interval between successive tones was 650ms, resulting in a stimulation frequency of 1.54Hz. Stimuli were thus comparable to those used in previous EEG studies on individual grouping (Brochard et al., 2003; Poudrier, 2020).

**Figure 1.**
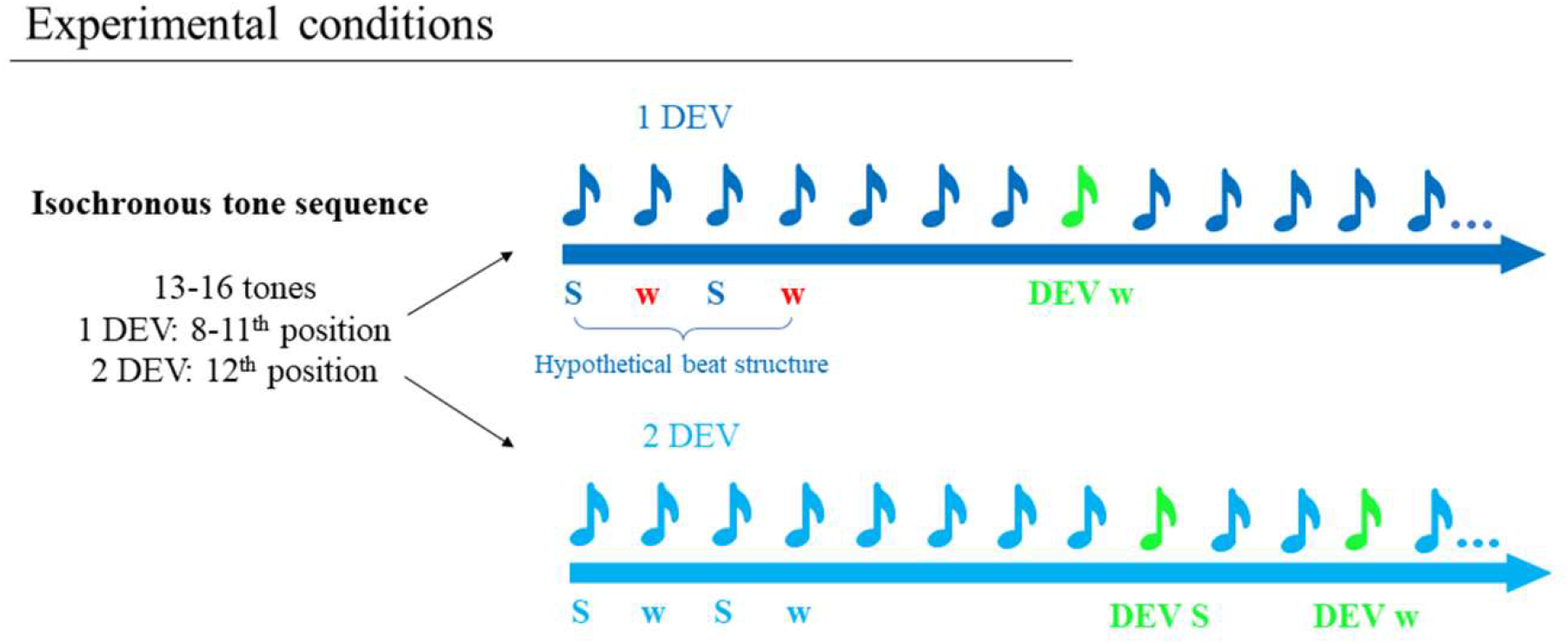
Experimental conditions. Participants listened to 192 isochronous tone sequences, containing 13-to-16 tones and either one or two deviants (DEV). The first DEV could either fall on positions 8,9,10,11^th^, while the second DEV always fell on position 12^th^. A hypothetical binary grouping structure could segment adjacent tones as Strong-weak (S-w; or on-/off-beat) tones. Thus, the first DEV could occur with equal probability in S-w tone positions.

Participants were seated in a dimly lit soundproof chamber facing a computer screen. A trial started with a fixation cross (500ms), followed by the presentation of the tone sequence. The cross was continuously displayed on the screen to prevent excessive eye movements while tone sequences were played. Immediately after the end of a sequence, a response screen appeared and prompted participants to press a response button to indicate whether they heard one or two softer tones. Button assignments were counterbalanced across participants. The inter-trial interval was 2000ms. A session was divided into two blocks of approximately 10 minutes each, with a short pause in between. Therefore, an experimental session lasted for about 25 minutes.

### 2.3. EEG recording

The EEG was recorded from 59 Ag/AgCl scalp electrodes (Electrocap International), amplified using a PORTI-32/MREFA amplifier (DC to 135 Hz), and digitized at 500 Hz. Electrode impedances were kept below 5kΩ. The left mastoid served as online reference. Additional vertical and horizontal electro-oculograms (EOGs) were recorded.

### 2.4. Data Analysis

#### 2.4.1. Behavioral analysis

Behavioral data (i.e.,response accuracy) were subjected to a repeated-measures ANOVA with deviant position (odd vs. even) as the independent variable and sequence order as a covariate.

#### 2.4.2. EEG Preprocessing

Data were pre-processed using combined custom Matlab scripts/functions and the Matlab-based FieldTrip toolbox (Oostenveld et al., 2011). Data were re-referenced to the average of the two mastoid electrodes, band-pass filtered with a 4th order Butterworth filter in the frequency range of 0.1-50 Hz (*ft_preprocessing*). Eye-blinks and other artifacts were identified using independent component analysis (‘*fastICA’* implemented in FieldTrip). A semi-automated pipeline was used to identify EEG components with a strong correlation (>.4) with the EOG timecourses, to inspect the respective topographical distribution across scalp electrodes and to remove “bad” components. Next, data segmentation was conducted separately for the rhythm-tracking analyses, event-related potential (ERP), and time-frequency representation (TFR) analyses.

#### 2.4.3. Rhythm tracking analyses

Rhythm-tracking analyses involved neural responses to the full auditory sequences. Following ICA, we created 192 (96 auditory sequences * 2 sessions per participant) 11s-long segments starting from the third tone onset up to the 13^th^ tone offset. Next, we created a fronto-central channel cluster, encompassing the sensor-level correspondents of prefrontal, pre-, para-, and post-central regions highlighted in previous studies (Fujioka et al., 2012; Fujioka et al., 2015). The cluster included 16 channels: ‘AFz’, ‘AF3’, ‘AF4’, ‘F3’, ‘F4’, ‘F5’, ‘F6’, ‘FCz’, ‘FC3’, ‘FC4’, ‘FC5’, ‘FC6’, ‘C1’, ‘C2’, ‘C3’, ‘C4’. Data from this fronto-central cluster were used for Fast-Fourier transform (FFT) and phase-locking analyses.

##### Fast-Fourier transform

Single-trial data from the fronto-central cluster were submitted to a FFT (referred to as FFT data) with an output frequency resolution of 0.14Hz. Spectral power was calculated as the squared absolute value of the complex Fourier output. Data in each frequency bin were normalized by the frequency-specific standard deviation across trials. Lastly, we averaged the frequency-domain data across channels and trials. For illustration purposes only, we restricted the Fourier spectrum from 1 –to 4Hz (Fig. 2A).

**Figure 2.**
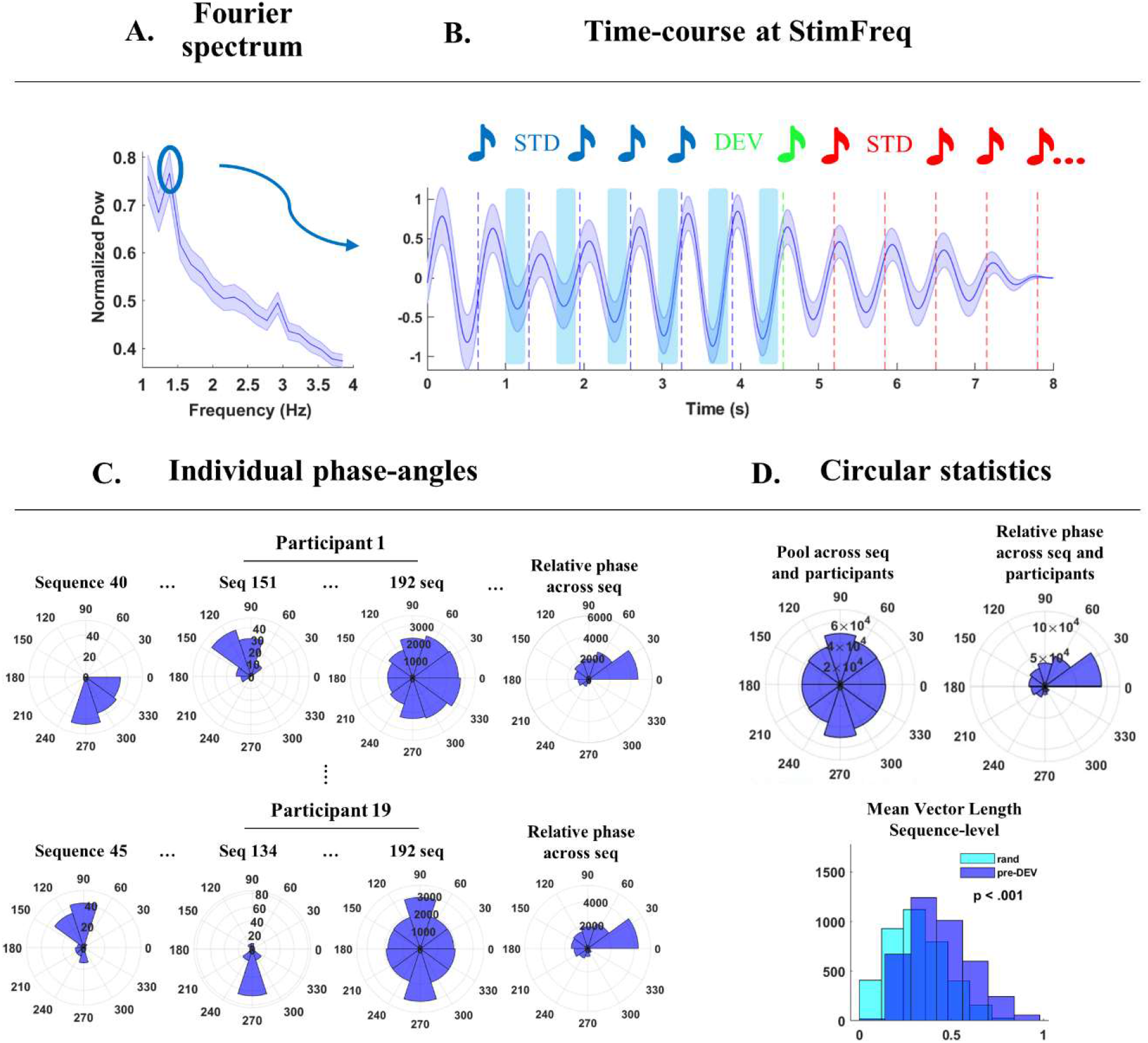
Rhythm tracking analyses A: Fourier spectrum of neural activity along the entire auditory sequence. The plot displays the grand-average normalized power in the frequency range from 1 to 4Hz. The blue circle highlights the peak amplitude for neural activity at the stimulation frequency (1.54Hz). B: time-course of neural activity at the stimulation frequency. Vertical bars indicate the onsets of STD tones prior-(blue) and post-DEV (red). The DEV onset is reported in green. Blue shades represent the standard errors. Light-blue rectangles indicate the focus on the pre-stimulus intervals (not scaled). C: Polar histograms for single-participant and sequence-level phase angles extracted from 60ms prior to the onsets of STD tones prior (blue) to the DEV from the fronto-central cluster of interest. Here, we report a few sequence-level phase-angles from Participant 1 (top) and Participant 19 (bottom). On the right, the polar histograms report the distribution of phase-angles across all trials (192 per participant) and the ‘relative phase’ across sequences. This is a measure of deviation from the most common phase-angle, at the sequence-level. D. Group-level phase-angles are randomly distributed around the polar histogram. On its right, the group-level ‘relative phase’. These phase-angles indicate a variation from the most common phase-angle. At the bottom, the distribution of mean vector length (MVL) calculated at the single-participant and sequence-level and averaged across the fronto-central cluster of interest. Importantly, these MVLs are based on the raw phase-angles for pre-DEV (blue) and are statistically compared to the MVL for random distribution of phase-angles. Single-participant statistics are reported in Suppl. Tab. 1.

##### Phase-locking analyses

A time-resolved phase-locking analysis was performed to estimate the phase relationship between neural activity at the stimulation frequency and the sequential tone onsets.

Full-sequence data from the fronto-central cluster were bandpass-filtered with a 4th order butterworth filter around the stimulation frequency (1.04-2.04Hz, to obtain a 1.54Hz center frequency; *ft_preprocessing*) and Hilbert-transformed to extract the analytic signal. Next, we plotted the timecourse of the real part of the analytic signal (Fig.2B) as a function of the STD tone onset preceding (blue) and following (red) the DEV (green; this plot is for illustrative purposes only). Phase-locking analyses were performed at the sequence and channel levels by means of circular statistics (circular toolbox in Matlab; (Berens, 2009)) and based on the circular mean phase-angles estimated in the ∼60ms (time-window proportional to the stimulation frequency = 1/1.54Hz/10) preceding individual tone onsets. Next, we calculated the sequence- and channel-level mean vector length (MVL; (Berens, 2009)) for pre-DEV STD tones and averaged the values across channels. MVLs for pre-DEV STD tones were statistically assessed against the MVL from a random distribution (random uniform distribution of phase-angles) by means of 1000 permutation tests. A p-value lower than .05 was considered statistically significant. For illustrative purposes, we also calculated participant-, channel- and sequence-level ‘relative phase angles’: these were expressed as the absolute phase difference between phase-angles for each tone position (e.g., 3^rd^ to 8th) and the most common phase-angle in the sequence. Examples of participant- and sequence-level relative phase-angles areplotted in Fig. 2C, and the pooling over participants, sequences, and channels is provided in Fig. 2D.

#### 2.4.4. ERP and TFR data

After ICA, data were segmented into 4s-long epochs symmetrically time-locked to every tone onset. Next, we employed an automatic channel-by-channel, trial- and participant-level artifact suppression procedure (similar to Kaneshiro et al., 2020). Amplitude values were temporarily normalized by their standard deviation across trials and outliers (data points per epoch and channel) were defined by means of a threshold criterion (values > mean + 4*SD). The identified noisy time-windows (with 50ms symmetrical padding) then served to suppress (replace by NaNs) time-points in the non-normalized data and these missing values were replaced by means of cubic temporal interpolation (*‘pchip’* option for both the built-in Matlab and FieldTrip-based interpolation functions) considering the time-course of neighboring time-windows (extending up to 700ms when possible, automatically reduced otherwise). Thus, rather than rejecting entire epochs containing artifacts (i.e., standard artifact rejection procedure), we opted for an artifact suppression approach that kept all trials (Kaneshiro et al., 2020). The channel-by-channel routine allowed the algorithm to flexibly adapt the outlier threshold estimates to the inherent noise varying over channels. On average, 20% of trials required the artifact suppression procedure. Lastly, a standard whole-trial rejection procedure based on an amplitude criterion (85uV) was applied. Data for event-related-potential (ERP) analyses (“ERP data”) were segmented, including 500ms prior and following each tone onset (1s in total). Data for the time-frequency representation analyses (“TFR data”) were not further segmented at this stage. ERP data were band-pass filtered between 1-30Hz and TFR data low-pass filtered at 40Hz. Data were then downsampled to 250Hz.

TFR data underwent time-frequency transformation by means of a wavelet-transform (Cohen, 2014), with a frequency resolution of .25Hz. The number of fitted cycles ranged from 3 for the low frequencies (<5Hz) to 10 for high frequencies (>5Hz and up to 40Hz). TFR data were then re-segmented to reduce the total length to 2s, symmetric around tone onsets.

##### Post-processing of ERP and TFR data

Single-trial ERP amplitudes were mean-corrected by a global average over all epochs (−0.2 to 0.3s relative to tone onset). Similarly, single-trial TFR power was normalized by computing relative percent change with reference to the global mean power across epochs (−0.2 to 0.3s relative to tone onset). This previously applied approach (Abbasi & Gross, 2020; Fujioka et al., 2012) was preferred over classical baseline correction because we aimed at analyzing power fluctuations in the pre-stimulus intervals. Finally, we calculated a fronto-central channel cluster average (using the same channels as for the rhythm tracking analyses). All subsequent analyses were performed exclusively on this channel cluster.

##### ERP analyses

Evoked responses over trials were averaged separately for STD and DEV tones, and for odd (hypothetical “Strong” position in a binary beat; S, Fig. 3A, left) and even (“weak”; w) serial positions from the 3rd to the 11^th^ tone. The first STD tones were disregarded to exclude the increased responses typically observed at the beginning of an auditory sequence. Fig. 3A shows the respective ERPs for the averaged fronto-central cluster, for STD tones in metrically strong and weak positions (left) and for the comparison of STD and DEV tones averaged over metric positions (right). Statistical analysis was performed by means of paired-sample t-tests. An adjusted p-value lower than .05 was considered statistically significant (Benjamini & Hochberg correction).

**Figure 3.**
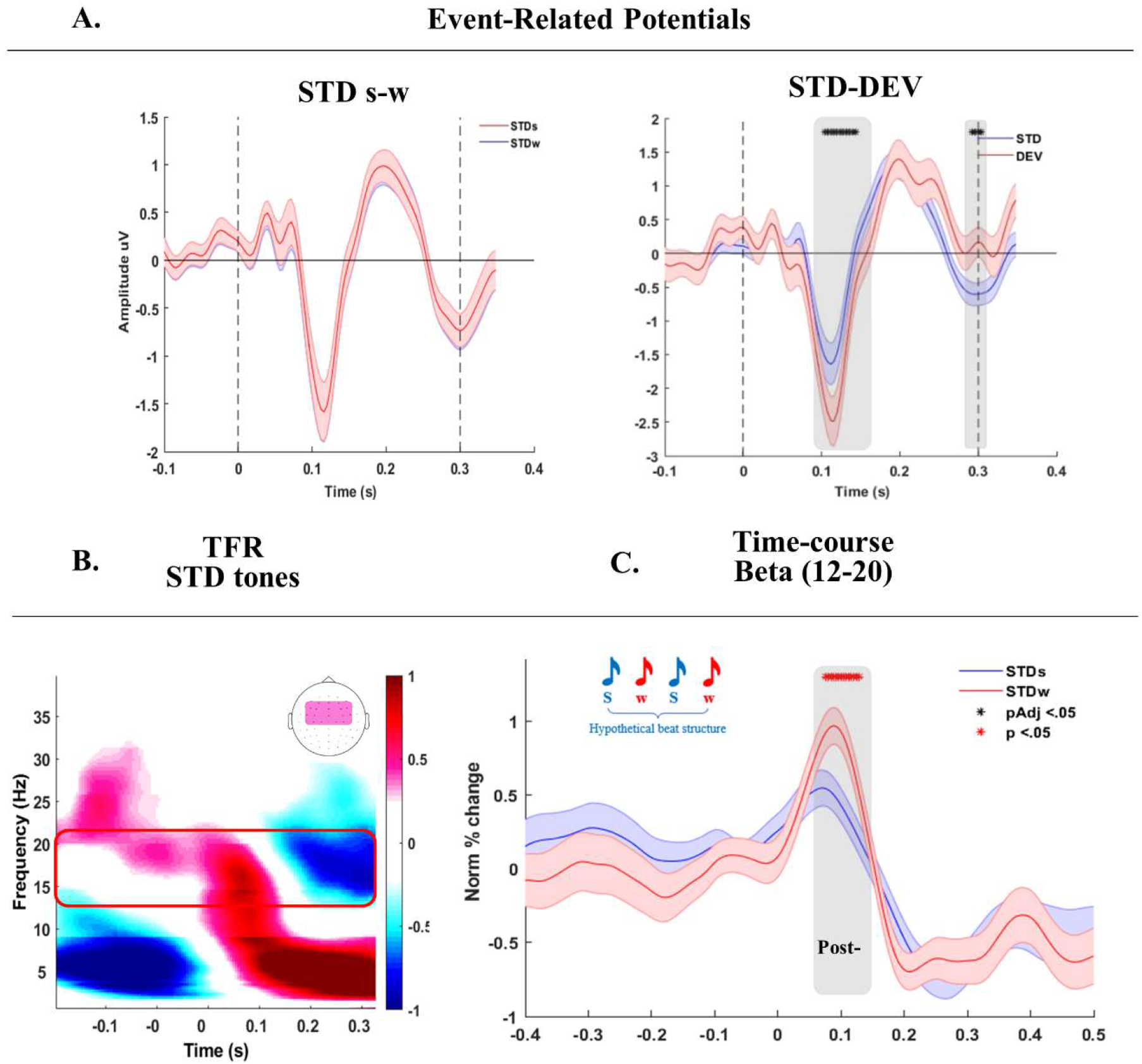
– Binary beat processing on hypothetical S-w positions. A: On the left, ERP responses for STD tones in S (blue) w (red) positions. On the right, ERP responses for STD (blue) and DEV tones. Stars indicate significant time-windows, as assessed by means of paired-sample t-tests (FDR-adjusted p <.05). B: grand-average time-frequency spectrum time-locked to STD tones (−.2 to .35s). The frequency range spans from 1-to-40 Hz with a frequency resolution of .25Hz. The red rectangle highlights evoked responses in the low-beta (12-20Hz) frequency range, on which we performed statistical comparisons in C. The topographic plot on top displays the FC cluster average in use. C: extracted time-course of low-beta activity in hypothetical S-w positions, time-locked to STD tones onsets, in blue for odd-numbered positions (Strong binary grouping) and red for even-numbered positions (weak binary grouping). Shaded colors report standard errors. On top, a grey rectangle delineates the time-window in which statistical testing showed a difference between S and w positions.

##### TFR analyses

Time–frequency representations were averaged over STD trials, separately for odd and even positions (hypothetical strong and weak positions, respectively; Fig. 3B). Next, we quantified mean amplitudes in the low-beta band (low-β; 12-20Hz) in the post-stimulus intervals (150ms after stimulus onset) and compared them for S-w positions (Fig. 3C).

##### Individual beat classification

Individual beat processing differences were assessed by means of a modelling approach suited to reveal binary, ternary, and potential other grouping patterns. We focused on single-participant, low-β power peak for STD tones in the first 8 positions of the auditory sequence in a 60ms time-window (proportional to the center frequency of interest; for low-β: 1/16Hz = 60ms) following the stimulus onset (from now on, β-post).

The beat modelling approach was based on β-post mean power, which was expected to reveal subjective accentuation patterns (Fujioka et al., 2012; Fujioka et al., 2015). Single-tone β-post were first concatenated to mimic an 8-tone auditory sequence (i.e., a trial). Single-participant and trial-level β-post (8 tones) were then entered into a stepwise regression model (Fig. 4A) (‘*stepwiselm*’ in Matlab) with 3 predictors: a binary (values: 1, −1), a ternary (1, −.5, −.5), and a constant term (ones). The winning model was chosen based on adjusted eta-squared. Trials for which the winning model involved the binary predictor were labeled “binary”, independent of whether the winning model included the intercept term. Similarly, trials for which the winning model involved the ternary predictor were labeled “ternary” independent of whether the intercept was included in the model. The model thus allowed the combination of multiple predictors, but no interactions between terms. Accordingly, we interpreted (and labeled) the combination of binary and ternary terms as “combined”. Last, trials in which grouping could not be clearly identified were labeled as “not classified”. Participant-level model results are provided in Suppl. Tab. 2 as “beat preferences” and expressed as the percentage of trials relative to the whole number of auditory sequences (192 per participant). The subject-level goodness of fit of the model is provided in the same table. The “Beat preferences” across participants are provided in Fig. 4B.

**Figure 4.**
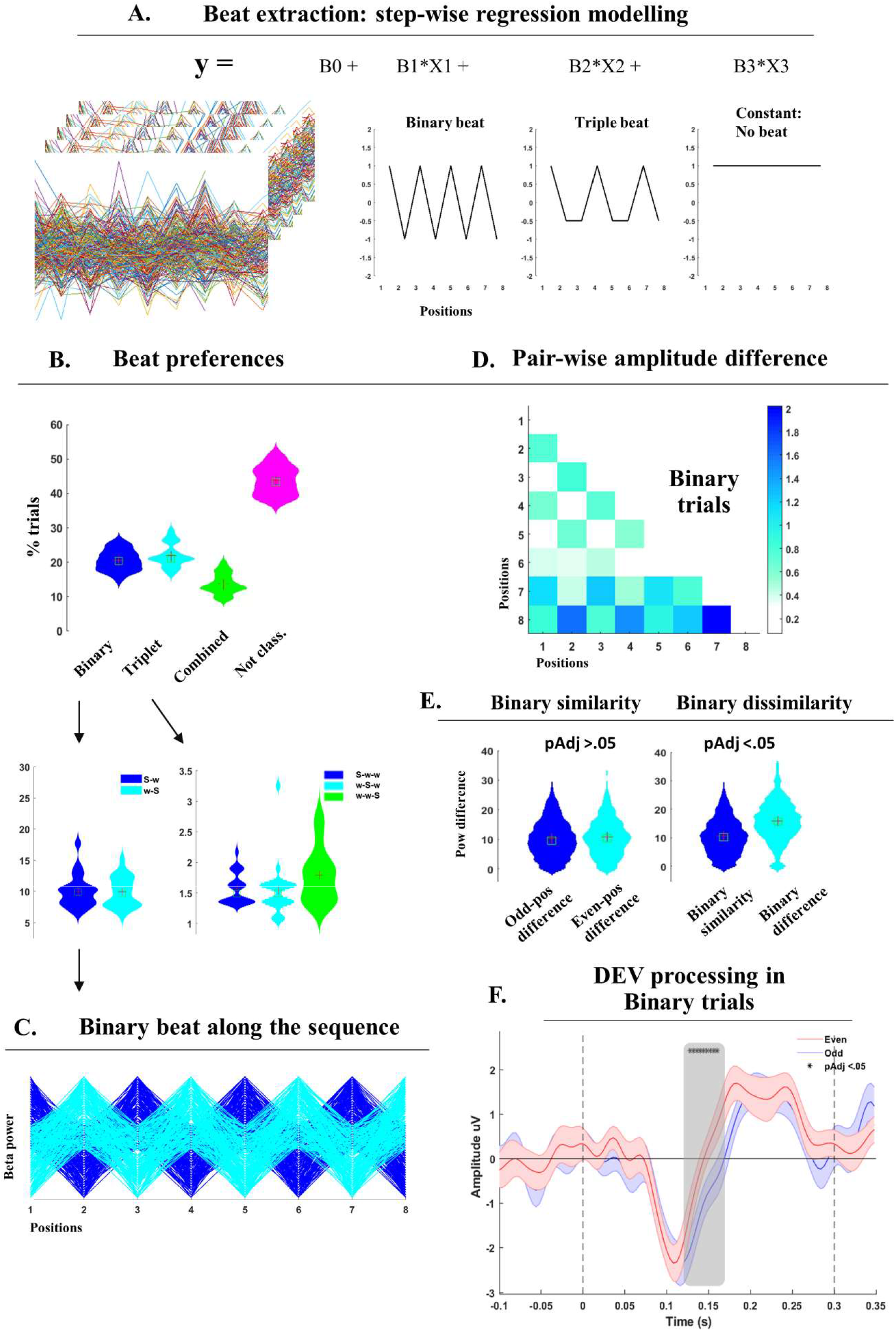
– Beat modelling and binary grouping. A: Beat modelling was performed by means of stepwise regression modelling and using low-beta post-stimulus responses as a dependent variable. The predictors were a binary (1, −1), a triple (1, −.5, −.5) and a constant term (ones). B: grouping preferences, as reported from the beat modelling. In order, we plot the distribution of trials assigned to binary, ternary, combined (binary-ternary) grouping, and ‘not classified’ (neither binary nor ternary) across participants. At the bottom, we zoom into binary trials and distinguish S-w accents from w-S accents based on trial-level Beta coefficients from the beat modelling. Similarly, on its right side, the distribution of ternary trials showing S-w-w, w-S-w, and w-w-S accentuation patterns. To extract these three accentuation patterns, we performed separate step-wise regression modelling as explained in the method section. C: Exemplar S-w and w-S accent fluctuations expected along 8 positions of the auditory sequence in the ‘binary’ grouping trials. Blue for S-w; cyan for w-S sequences. This plot has illustration purposes only. D: grand-average pair-wise difference for low-beta peaks across the first 8 positions of the auditory sequence in binary trials. E: on the left, the distribution of amplitude differences across odd-numbered positions (in blue) and even-numbered positions (cyan). The average of these two distributions forms the ‘Binary similarity’. On the right, the ‘binary similarity’ (blue) and the mean amplitude difference of odd-versus even-numbered position (‘binary dissimilarity’; in cyan). Statistical testing was performed by means of thousand permutation testing, and an FDR-adjusted p < .05 was considered as statistically significant. F: ERPs to DEV tones on S-w positions in the binary trials. Statistical testing reported a significant difference in the time-window between ∼120-170ms post-stimulus, as highlighted by the grey shades.

##### Binary group analyses

Further analyses were performed to verify if the identified “binary” trials indeed showed binary-like accentuation patterns. If so, neural responses to tones falling on odd-numbered positions should differ from those on even-numbered positions. However, there should be no differences for neural responses on the same metrical positions: namely, tones falling on odd-numbered positions should elicit similar (i.e., non-significantly different) neural activity.

To test these hypotheses, we first vertically concatenated trials classified as “binary”, and then computed a trial-based single-tone pair-wise amplitude difference (corresponding lower-triangle 2-D means are provided in Fig. 4D), i.e., amplitude differences between responses to each tone in the sequence (1-8 positions). For example, the response amplitude for the 1^st^ position was compared to the 2^nd^ position, then to the 3rd, and so forth. In turn, amplitude for the 2^nd^ position was compared to the 3^rd^, the 4^th^etc. The resulting pair-wise amplitude difference matrix had a size of N trials x N positions-1 x N positions-1. From this matrix, we statistically compared the pair-wise amplitude difference for tones in odd-positions (Fig. 4E; “odd-pos difference”) and even-positions (“even-pos difference”) by means of 1000 permutations of odd-even labels. An FDR-adjusted p-value lower than .05 was considered statistically significant (Benjamini & Hochberg correction). The two variables were then combined into a distribution of “binary similarity”. The binary similarity combines the amplitude difference for tones in odd-numbered positions (1-3-5-7^th^) and the amplitude difference for tones on even-numbered positions (2-4-6-8^th^). Binary similarity was statistically compared to “binary dissimilarity”, which was calculated as the mean difference of tones in odd versus even positions (Fig. 4E). Statistical testing was performed using 1000 permutations of odd-even labels, and an FDR-adjusted p-value lower than .05 was considered statistically significant (Benjamini & Hochberg correction).

##### DEV analyses as a function of binary beat

For each participant, we isolated binary trials identified with the described “single-participant beat classification” procedure to explore the relation of beat and DEV processing as indicated by a differentiation of ERP responses according to their accents (S-w). Statistical analyses compared ERPs to DEV tones in even- (8,10th) versus odd-numbered (9,11th) positions (Fig. 4F). Notably, we considered possible “pure binary” and “phase-shifted binary” beats (or inverse binary), where the former grouping corresponds to the typical S-w pattern in odd and even positions, and the second the reverse w-S pattern. Indeed, individuals may start grouping at different times along the auditory sequences, and hypothetical S-w positions may likely fall on either odd or even positions. The selection was informed by the beta coefficients associated with the identified binary trials: a positive coefficient would indicate “pure binary” (S-w), while a negative coefficient would correspond to the “inverse binary” (w-S). The distribution of “pure” and “inverse” binary is provided at the bottom left of Fig. 4B, expressed as percent of trials relative to the total number of auditory sequences (192 per participant).

Next, we re-ordered accented S-w positions according to individual binary processing (odd-numbered positions falling on S beats in “pure binary” but on w in “inverse binary”), and statistically compared their associated ERPs time-courses (Fig. 4F). Statistical testing was performed by means of paired-sample t-tests, and an FDR-adjusted p-value lower than .05 was considered as statistically significant (Benjamini & Hochberg correction).

##### DEV analyses in non-classified trials

We also tested whether a similar S-w effect would be observed for DEV processing in ‘non-classified’ trials. Evoked responses for DEV tones falling on accented S (odd-numbered positions) and w (even-numbered) positions were pooled and statistically compared by means of paired-sample t-tests. An FDR-adjusted p-value lower than .05 was considered as the accent can either fall on the first (S-w-w), second (w-S-w), or third (w-w-S) positionHowever, with, we could not adopt the same procedure. In order to disentangle these three accentuation patterns from the distribution of “ternary beats” identified during the “single-participant beat classification” procedure, we ran a second step-wise regression model. The model featured three predictors mirroring the above-mentioned accentuation types, and whose values were respectively: 1,−.5,−.5 (pattern 1), −.5,1,−.5 (pattern 2) and −.5,−.5,1 (pattern 3). The model did not allow interaction terms, and the winning model was chosen based on adjusted eta-squared. The output of the model is provided in Fig. 4B (bottom right), as the percentage distribution of three accentuation patterns across participants

## Acknowledgements

We thank Ina Koch at the Max Planck for Human Cognitive and Brain Sciences, Dept. of Neuropsychology, Leipzig, Germany for her support in data collection. This work is supported by a grant of the Dutch Research Council (NWO - 406.18.GO.063) and the Van der Gaag Fund, Royal Netherlands Academy of Arts & Sciences.

## 3. Code Accessibility

The code in use for analyzing this dataset has been stored and can be provided upon reasonable request by the corresponding author.

## 4. Results

### 4.1. Behavioral Data

We tested whether the counting of deviant tones (DEV) differed for deviants in odd or even positions in the auditory sequence. The respective ANOVA with deviant position (odd vs. even) and sequence order as a covariate did not result in a significant effect of deviant position (F(1,71) = 1.115, p = .295, eta-square = 0.16) nor a significant effect of sequence order (F(1,69) = .02, p = .97, eta-square = 0). Thus, DEV counting performance did not differ in the auditory sequence.

### 4.2. Rhythm tracking

Participants listened to isochronous auditory sequences presented at a stimulation rate of 1.54Hz and we tested *whether* and *how* their neural activity would show idiosyncratic signatures of rhythm tracking.

When individuals listen to these sequences, their neural activity reflects the timing of external events (Fig.2A-B). Indeed, the Fourier spectrum showed a clear peak at the stimulation frequency (1.5Hz; Fig. 2A) and the time-course of delta-band neural activity showed a tendency to align to tone onsets (Fig. 2B). To quantify the consistency of anticipatory phase alignment to the expected tone onsets, we tested the phase consistency of delta-band (1.5Hz) neural activity in a time-window preceding tones onset. Thus, phase-locking analyses focused on the ∼60ms (proportional to the stimulation frequency: 1/1.5Hz/10) prior to tone onsets. Single-participant trial-level mean vector lengths (MVL) of STD tones preceding a DEV (pre-DEV) revealed a consistent phase-relationship with STD onsets: the MVL significantly differed from a random distribution (Fig. 2D, bottom; pre-DEV in blue; Suppl. Tab. 1 for statistical results). Importantly, however, we observed intra- and inter-individual differences: participants’ delta-band activity did not always synchronize to tone onsets with the same phase relationship (Fig. 2C). Rather, a broad range of possible phase-lags was observed across trials, both at the level of single-participants (Fig. 2C right) as well as when pooling values across participants (Fig. 2D top-left). Hence, the distribution of single-participant phase-angles across trials did not differ from a random distribution (Suppl. Tab. 1). Phase-angles were consistent within a trial (MVL statistics in Fig.

2D bottom and Suppl. Tab. 1), but differed across trials. To further explore this variability, we computed a measure of ‘relative phase’. This was calculated, at the single-participant, channel- and sequence-level as the absolute difference from each phase-angles within the sequence and the most common phase. The distribution of relative phase-angles across trials and participants shows a variance which mostly ranges between 0-30 degrees (Fig. 2D, right), supporting the MVL calculation. Thus, individuals show a predictive and consistent phase-alignment of delta-band activity to expected tone onsets. However, the specific phase for alignment is variable across trials.

### 4.3. Beat processing

We tested whether participants’ neural activity would show signatures of spontaneous beat processing, visible as binary-like accentuation patterns (S-w accents in odd-numbered versus even-numbered positions). Thus, we tested event-related responses (ERP) to STD tones in S and w positions, and further inspected the time-frequency representation of time-locked responses. ERPs to STD tones in S-w positions did not statistically differ (Fig. 3A). However, DEV tones elicited stronger N100 responses as compared to STD tones (FDR-adjusted *p* < .05; Fig. 3A, right), confirming the processing of an unpredicted deviant tone.

The time-frequency representation plots of neural activity in response to STD tones mainly showed two event-locked responses (Fig. 3B): one in the theta (4-8Hz) and one in the low-beta (low-β; 12-20Hz) frequency-band. Following our hypotheses, we focused on the time-course of activity in the low-β range and compared event-locked power fluctuations for STD tones on odd (S) to even (w) numbered positions along the sequence (Fig. 3C), corresponding to hypothetical S-w binary grouping (blue and red, respectively). Thus, we statistically compared the time-courses of low-β activity (Fig. 3C) for STD tones in hypothetical S-w positions. Statistical testing did not survive multiple comparison by FDR correction.

In summary, neither ERP nor TFR analyses revealed a binary beat effect, as STD tones elicited similar responses when they occurred in odd- and even-numbered positions in the auditory sequence. We hypothesized that this result may be associated with inter- and intra-individual differences in *when* and *how* accentuation patterns are superimposed onto the auditory sequences. Namely, individuals may start to accentuate at different points along the sequence (i.e., not necessarily at the beginning), may do it differently over time (e.g., binary or ternary accents), or may even not accentuate at all (Brochard et al., 2003). These hypotheses were tested with the novel beat modelling approach below.

### 4.4. Beat modelling

To address the questions of (i) whether everyone accentuates in a similar way, (ii) whether everyone always accentuates in the first place, and (iii) whether accentuation patterns influence DEV processing (Brochard et al., 2003), we focused on trial-level data and modelled various accentuation patterns. In particular, we used a trial-based stepwise regression model to classify participant-level single-trial beta-band neural responses as best reflecting a binary, ternary, combined, or ambiguous grouping. The choice of beta-band activity was informed by previous evidence linking it to beat processing (Fujioka et al., 2012; Fujioka et al., 2015). The model predicted tone-by-tone low-β-post peak power from three predictors: binary, ternary, and constant terms (Fig. 3A). Resulting grouping preferences and goodness of fit are reported in Suppl. Tab. 2 and summarized in Fig. 4B.

Furthermore, we zoomed into “binary trials”, and distinguished S-w from w-S accentuation patterns based on the single-trial β-coefficients from the beat modelling (see methods). The resulting distributions are reported in Fig. 4B, bottom left. Similarly, we disentangled three possible accentuation patterns in the “ternary trials”. We performed a separate step-wise regression model using S-w-w, w-S-w, and w-w-S accents as predictors (see methods). Distributions are reported in Fig. 4B, bottom right.

This approach allowed showing that individuals spontaneously superimpose accentuation patterns on identical tones embedded in an isochronous equitone sequence. Importantly, we confirm that participants switch between binary, ternary, and other accentuation patterns over trials. However, in the majority of trials no consistent accentuation pattern was confirmed.

### 4.5. Binary beat

Once we isolated, at the single-participant level, trials showing binary-like accentuation patterns, we aimed to statistically test whether low-β responses would significantly differ in S versus w positions. Thus, we tested whether the beat modelling approach delivers a meaningful classification of binary grouping.

We isolated the identified ‘binary’ beat trials and calculated the tone-by-tone pair-wise difference for low-β across 8 positions in the auditory sequence and preceding the DEV tone. For visualization purposes, the resulting matrix was averaged across trials and the upper symmetrical triangle was masked (Fig.4D). The original matrix (all trials) was used to calculate metrics of “Binary similarity” and “Binary dissimilarity” (Fig.4E; see ‘Binary group analyses’ in the methods). The Binary similarity features the distributions of amplitude differences on odd- and even-numbered positions. For the “Binary dissimilarity” analyses we calculated the amplitude difference for tones on odd-versus even-numbered positions (corresponding to on-beat versus off-beat; thus labeled “Binary difference”) and statistically compared it to the Binary similarity (right-side plot in Fig.4E). Statistical testing yielded a significant difference (FDR-adjusted *p* < .05).

In summary, we confirmed that the trials classified as ‘binary’ in the beat modelling, do indeed show a consistent binary accentuation patter. Hence, the low-β amplitudes in STD tones in S positions significantly differ from those in w positions. To further verify the validity of the beat modelling approach, we tested whether identified ‘beat preferences’ modulate DEV processing.

### 4.6. DEV processing based on binary grouping

We investigated whether DEV processing is modulated by a binary beat in ‘binary’ beat trials. Thus, we tested whether ERPs to DEV tones falling on S-w beat positions in the identified “binary” trials would be statistically different. First, we isolated the identified binary trials and discerned ‘pure binary’ from ‘inverse binary’ trials based on the beta-coefficient resulting from the regression modelling (see methods). Next, we pooled together odd-numbered (9,11^th^ positions) and even-numbered (8,10^th^) DEV positions (i.e., corresponding to S-w). Within-participant statistical comparison of the respective ERPs yielded significant difference in the time-window between 120-170ms (*p FDR adjusted <*.*05*; Fig. 4F).

Similarly, we tested whether the same S-w effect would be observed for those trials in which no accentuation pattern could be identified (‘non-classified’ trials). In these non-classified trials, DEV processing was not modulated by a binary beat (*p FDR adjusted >*.*05*; Suppl. Fig. 1).

Lastly, we focused on the ‘ternary’ trials and modelled three possible accentuation patterns: S-w-w, w-S-w, w-w-S (Fig. 4b). Given the small percentage of trials belonging to the three accentuation types (∼2%), we refrained from performing further analyses due to insufficient statistical power to interpret results.

In summary, we here show that DEV processing is modulated by a binary beat, but exclusively in those trials identified as ‘binary’ during the beat modelling. This observation supports the beat modelling procedure as a viable method for identification of trial-level variability in beat processing.

## 5. Discussion

The current study aimed at exploring individual neurophysiological variability in rhythm and basic beat processing. In particular, we focused on rhythm-tracking by analysing the sequence-level phase-consistency of delta-band neural activity while predictively synchronizing to tone onsets. Next, we tested whether neural activity in the low-beta band (12-20Hz) would reflect the emergence of binary-like accentuation patterns, indicating spontaneous engagement in beat processing.

When listening to isochronous equitone sequences, participants’ neural activity tracked the timing of external events (Fig.2A), aligning delta-band oscillatory dynamics to expected tone onsets (Fig.2) (Buzsáki, 2009; Schroeder & Lakatos, 2009; Thut et al., 2012b; Zoefel & VanRullen, 2016). Hence, sequence-level mean vector lengths of delta-band activity preceding tone onsets displayed anticipatory coupling of brain activity to the timing of environmental stimuli. This finding further confirms theoretical views according to which the brain might generate temporal predictions to achieve successful rhythm tracking to optimize sensory processing, perception, and allocation of attention (Friston, 2005; Arnal, 2012; Schröger et al., 2015; Koelsch et al., 2019).

Notably, when pooling phase-angles across sequences either at the single-participant or group-level, we observed random phase distributions. In other words, delta-band neural activity did not always align its high-excitability phase to the expected onset of auditory tones. Rather, we observed a wide range of possible synchronization regimes, which show consistency at the sequence-level, but high variability across sequences and participants. These findings counter predominant views on entrainment that propose a specific phase-alignment (high-excitability phase) with environmental stimuli to optimize stimulus processing. The current findings rather suggest that entrainment may not necessarily rely on a fixed synchronization regime, but rather on flexible and adaptive phase-alignment, that can vary over time and across individuals, while keeping the consistency at the trial-level. However, task and attention manipulations may modulate the observed trial-level variability.

Next to rhythm tracking, we investigated the neural signatures associated with the human disposition to group two or three adjacent tones when listening to isochronous equitone sequences (Brochard et al., 2003). This spontaneous grouping induces binary (strong-weak (S-w)) or ternary (S-w-w) accentuations patterns (Brochard et al., 2003), and might resembles the superimposition of a beat structure. Importantly, while the tones are physically identical, the beat can influence observable behaviour and underlying neural activity (Nozaradan, 2014; Nozaradan et al., 2011, 2016, 2017; Schmidt-Kassow et al., 2011). So far, however, behavioural and neuroimaging research has dedicated little space to investigations of individual differences (Grahn & Brett, 2007; Grahn & McAuley, 2009) and mainly tested task-based beat processing (e.g., Fujioka et al., 2015). Thus, the question remains whether participants would naturally engage in beat processing in the absence of specific task instructions, and if they would do so in a similar manner. To address this question, we focused on participant-level neurophysiological variability and modelled single-trial beta-band activity (Fig.4A) to probe whether accentuation patterns mirror binary, ternary, and other forms of grouping. Neural activity in the beta range has been associated with rhythm (Arnal, 2012; Biau & Kotz, 2018; Fujioka et al., 2012; Fujioka et al., 2015; Morillon et al., 2016) and beat processing (Fujioka et al., 2012; Fujioka et al., 2010). The current findings confirm its prevalence in time-locked responses to STD tones (Fig.3B). Furthermore, beta-band activity showed an individuals’ spontaneous disposition to superimpose a beat pattern, even when not instructed to do so (Fig.4). Hence, we characterized inter- and within-participant differences in adopting binary, ternary, and combined grouping over time (Fig. 4B and Suppl. Tab. 2). Beat processing in the absence of specific task instruction might reflect the automatic predisposition to temporally organize continuous auditory input streams into predictable, coherent, and finite units (i.e., beat groups). This might reflect fluctuations of attentional resources over time (Bolton, 1984; Jones, 1976; Schroeder & Lakatos, 2009) (rather than being equally distributed over time), and variations within each attentional cycle, so to attribute salience to on-beat events. We tested this view by particularly focusing on binary grouping (Fig.4C-F) and showed that beta amplitudes of tones falling on odd-numbered (on-beat or ‘strong’ (‘S’)) positions were significantly different from those in even-numbered positions (off-beat or ‘weak’ (‘w’); Fig. 4E,F). This suggests that individuals superimpose binary grouping (S-w beat) while listening to auditory sequences. Such grouping resembles the active parsing and segmentation of continuous sensory streams and might serve to allocate attentional resources to salient sensory events to ultimately optimize perception (Nobre & Van Ede, 2018; Shalev et al., 2019).

We acknowledge that classifying trials based on beta fluctuations and then testing beta time-locked responses over sequence positions might be circular, but confirmatory. Thus, to test this further, we compared neural responses to unpredicted (deviant) tones falling on S-w positions and found a significant effect in ERP responses to DEV tones (Fig. 4F). Notably, the modulation of DEV processing was absent in those trials which did not adhere to a specific accentuation pattern (non-classified trials; Suppl. Fig. 1). These results parallel earlier results (Brochard et al., 2003; Jongsma et al., 2004; Schmidt-Kassow et al., 2011) and indicate that beat processing might affect how we allocate attention to the auditory environment. However, further investigations are needed to clarify this intricate link between attention deployment, attentional shifts, and beat processing in listening contexts.

In summary, we identify individual rhythm tracking and beat processing differences, and associate them with specific delta-band phase-coupling mechanisms and with beta-band dynamics, respectively. The findings showcase the feasibility of using EEG to identify individual neurophysiological signatures in rhythm cognition, suggesting that common trial- and group-level averaging approaches might inevitably obscure inter-individual differences and trial-by-trial variability. In contrast, the approach adopted here allows the monitoring of neurophysiological variability underlying *flexible but consistent mechanisms* for evaluating and adapting to (un)predictable environmental stimuli. Consequently, we propose that zooming into individual variability might allow to better predict behavioral variability in processing simple and complex environmental rhythms (e.g., speech tracking) in neurotypical and pathological populations (Schwartze et al., 2015, 2016).

## 6. Conclusions

When listening to isochronous equitone sequences, humans’ neural activity tends to spontaneously align and track the timing of auditory events. Moreover, the current results indicate that individuals can endogenously engage in beat processing by accentuating auditory sequences with binary, ternary, and combined beat patterns. We explored inter-individual and trial-level neurophysiological variability in rhythm cognition, and reveal flexible, time-varying neural mechanisms to effectively evaluate and adapt to rhythms of the external environment. The combined findings highlight that an individualized analysis approach to neurophysiological data can indicate meaningful variation in a listening context and should be considered in a more differentiated account of timing in audition.

## Supplementary materials

**Suppl. Table 1.** Circular statistics for phase-angles and mean vector lengths Statistical testing of the phase-angle differences between neural activity at the stimulation frequency (1.5Hz) for, in order, STD tones pre-DEV versus a random distribution and pre-DEV versus post-DEV. On the right, the statistical comparison of trial-based mean vector lengths (MVL) for STD tones pre-DEV versus a random distribution. The phase-angles are extracted, per tone, in the pre-stimulus intervals in a time-window of 60ms (1/1.5Hz/10) and averaged with circular mean statistics. The MVL are calculated based on the trial-level phase-angle circular means. Statistical testing was performed, independently, by means of 1000 permutation tests. In the table are provided, in order the p-values (rounded to 3 decimals), observed differences and effect sizes per subject.

|  | *Phase-angles*<br>*Pre-DEV Vs Random* |  |  | *Phase-angles*<br>*Pre-DEV Vs post-DEV* |  |  | *MVL*<br>*Pre-DEV Vs Random* |  |  |
| --- | --- | --- | --- | --- | --- | --- | --- | --- | --- |
|  | P vals | Obs Diff | Eff Size | P vals | Obs Diff | Eff Size | P vals | Obs Diff | Eff Size |
| **Subj1** | 0.233 | 1.733 | 1.277 | 0.057 | -2.715 | -2.039 | 0.001 | 0.145 | 0.726 |
| **Subj2** | 0.689 | 0.719 | 0.520 | 0.906 | 0.073 | 0.056 | 0.001 | 0.121 | 0.654 |
| **Subj3** | 0.505 | 0.917 | 0.671 | 0.336 | 0.694 | 0.523 | 0.001 | 0.110 | 0.593 |
| **Subj4** | 0.461 | -0.727 | -0.539 | 0.755 | 0.160 | 0.121 | 0.001 | 0.156 | 0.834 |
| **Subj5** | 0.261 | -1.629 | -1.195 | 0.740 | -0.176 | -0.136 | 0.001 | 0.197 | 1.099 |
| **Subj6** | 0.479 | 0.598 | 0.446 | 0.692 | 0.169 | 0.132 | 0.001 | 0.138 | 0.729 |
| **Subj7** | 0.805 | 0.230 | 0.171 | 0.859 | -0.089 | -0.068 | 0.001 | 0.085 | 0.469 |
| **Subj8** | 0.662 | -1.060 | -0.758 | 0.069 | -2.956 | -2.141 | 0.001 | 0.096 | 0.479 |
| **Subj9** | 0.358 | 1.818 | 1.315 | 0.149 | 2.448 | 1.776 | 0.001 | 0.113 | 0.607 |
| **Subj10** | 0.606 | -0.756 | -0.551 | 0.719 | -0.259 | -0.196 | 0.001 | 0.109 | 0.616 |
| **Subj11** | 0.802 | -0.360 | -0.262 | 0.919 | -0.100 | -0.075 | 0.001 | 0.124 | 0.655 |
| **Subj12** | 0.656 | 0.800 | 0.581 | 0.953 | 0.069 | 0.051 | 0.001 | 0.115 | 0.646 |
| **Subj13** | 0.057 | 2.993 | 2.170 | 0.059 | 2.962 | 2.151 | 0.001 | 0.089 | 0.474 |
| **Subj14** | 0.094 | 2.838 | 2.048 | 0.146 | 2.633 | 1.900 | 0.001 | 0.141 | 0.721 |
| **Subj15** | 0.093 | -2.489 | -1.836 | 0.582 | 0.236 | 0.186 | 0.001 | 0.107 | 0.556 |
| **Subj16** | 0.123 | -2.647 | -1.911 | 0.905 | 0.155 | 0.112 | 0.001 | 0.135 | 0.704 |
| **Subj17** | 0.583 | -1.229 | -0.881 | 0.973 | 0.056 | 0.040 | 0.001 | 0.110 | 0.605 |
| **Subj18** | 0.402 | 1.909 | 1.358 | 0.034 | -3.076 | -2.205 | 0.001 | 0.135 | 0.718 |
| **Subj19** | 0.778 | -0.704 | -0.504 | 0.339 | 1.127 | 0.839 | 0.001 | 0.122 | 0.643 |
| **Subj20** | 0.314 | 1.407 | 1.033 | 0.475 | -0.559 | -0.416 | 0.001 | 0.174 | 0.918 |

**Table 2.** Grouping preferences from the beat modelling. The table reports the percentages of individual preferences for (in order) binary, ternary, combined (binary + ternary) beat. The last column includes all trials which could not be classified as the above-mentioned beat strategies. The last column on the right provides the goodness of fit (expressed as Eta-squared, ‘R2’) of the chosen models, averaged across trials per subject.

|  | *Binary %* | *Ternary %* | *Combined %* | *Other %* | *R2* |
| --- | --- | --- | --- | --- | --- |
| **Subj1** | 19,3 | 26 | 9,4 | 45,3 | 0,40 |
| **Subj2** | 20,8 | 18,2 | 18,2 | 42,8 | 0,38 |
| **Subj3** | 20,3 | 22,9 | 16,1 | 40,7 | 0,39 |
| **Subj4** | 20,3 | 29,2 | 12,5 | 38 | 0,39 |
| **Subj5** | 22,4 | 21,4 | 17,7 | 38,5 | 0,41 |
| **Subj6** | 17,2 | 19,3 | 13 | 50,5 | 0,41 |
| **Subj7** | 18,2 | 26 | 12,5 | 43,3 | 0,37 |
| **Subj8** | 26 | 21,4 | 9,4 | 43,2 | 0,39 |
| **Subj9** | 17,2 | 20,8 | 14,1 | 47,9 | 0,36 |
| **Subj10** | 15,1 | 26,6 | 12 | 46,3 | 0,36 |
| **Subj11** | 22,4 | 17,7 | 13,5 | 46,4 | 0,37 |
| **Subj12** | 25 | 21,4 | 14,6 | 39 | 0,39 |
| **Subj13** | 17,7 | 20,8 | 15,6 | 45,9 | 0,39 |
| **Subj14** | 24 | 20,8 | 17,2 | 38 | 0,39 |
| **Subj15** | 17,2 | 20,8 | 19,8 | 42,2 | 0,38 |
| **Subj16** | 21,9 | 27,6 | 9,9 | 40,6 | 0,35 |
| **Subj17** | 22,9 | 18,8 | 10,9 | 47,4 | 0,34 |
| **Subj18** | 25 | 22,9 | 12,5 | 39,6 | 0,39 |
| **Subj19** | 19,8 | 16,7 | 12,5 | 51 | 0,36 |
| **Subj20** | 18,8 | 20,8 | 12 | 48,4 | 0,40 |

**Suppl. Figure 1.**
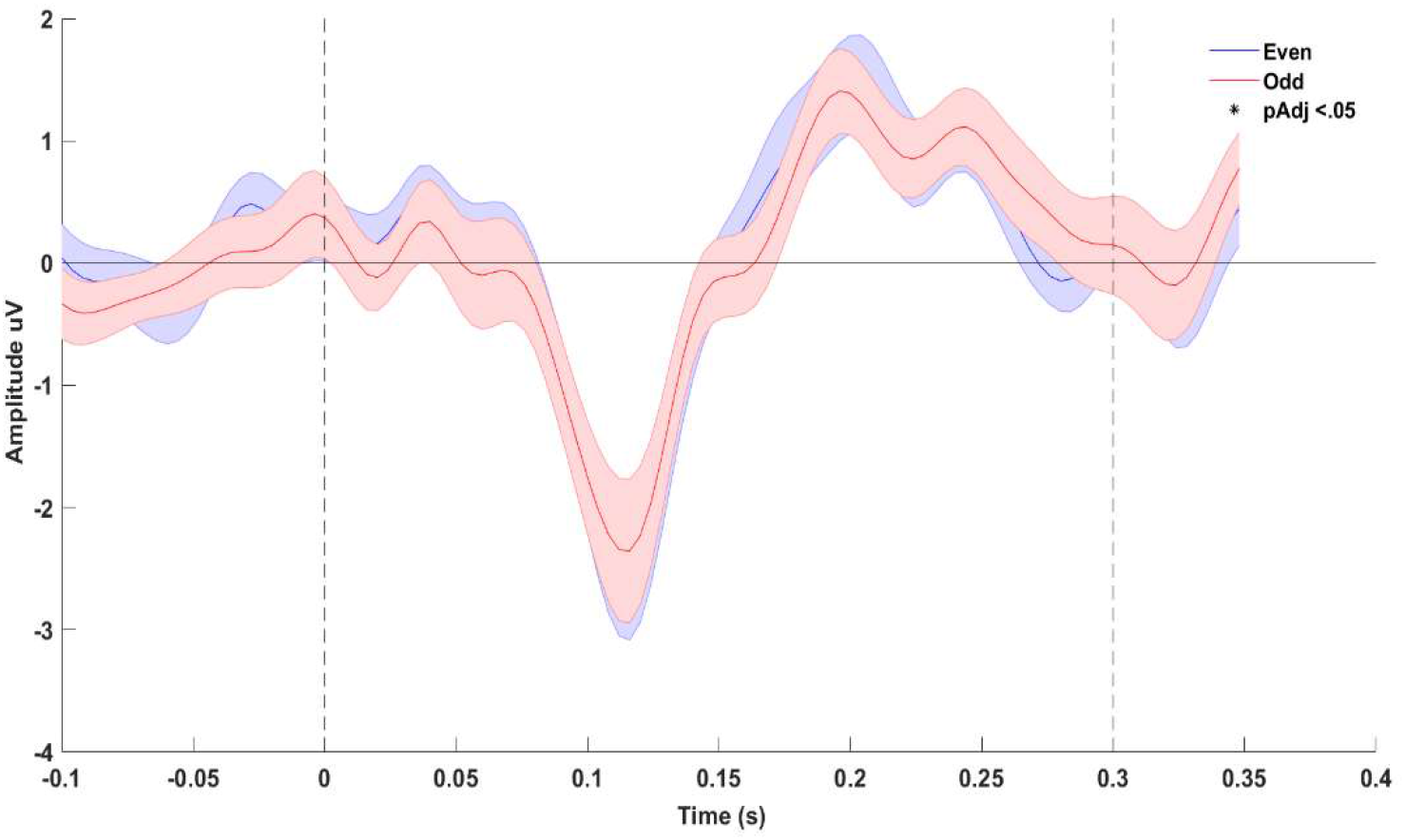
DEV processing is not modulated by a binary beat in non-classified trials Event-related responses to DEV tones falling on even-numbered (blue) and odd-numbered (red) positions along the auditory sequence and in those trials in which no accentuation pattern was observed (non-classified trials). Statistical comparison did not yield any significant difference.

## References

Abbasi, O., & Gross, J. (2020). Beta-band oscillations play an essential role in motor–auditory interactions. Human Brain Mapping, 41(3), 656–665. https://doi.org/10.1002/hbm.24830

Abecasis, D., Brochard, R., Granot, R., & Drake, C. (2005). Differential brain response to metrical accents in isochronous auditory sequences. Music Perception. https://doi.org/10.1525/mp.2005.22.3.549

Arnal, L. H. (2012). Predicting “When” Using the Motor System’s Beta-Band Oscillations. Frontiers in Human Neuroscience, 6. https://doi.org/10.3389/fnhum.2012.00225

Baath, R. (2015). SUBJECTIVE RHYTHMIZATION:A REPLICATION AND AN ASSESSMENT OF TWO THEORETICAL EXPLANATIONS. 244–254.

Barnes, R., & Jones, M. R. (2000). Expectancy, Attention, and Time. Cognitive Psychology. https://doi.org/10.1006/cogp.2000.0738

Bartolo, R., & Merchant, H. (2015). β oscillations are linked to the initiation of sensory-cued movement sequences and the internal guidance of regular tapping in the monkey. Journal of Neuroscience. https://doi.org/10.1523/JNEUROSCI.4570-14.2015

Berens, P. (2009). CircStat: A MATLAB Toolbox for Circular Statistics. Journal of Statistical Software, 31(10), 1–21. https://doi.org/10.18637/JSS.V031.I10

Biau, E., & Kotz, S. A. (2018). Lower beta: A central coordinator of temporal prediction in multimodal speech. Frontiers in Human Neuroscience, 12(October), 1–12. https://doi.org/10.3389/fnhum.2018.00434

Bolton. (1984). Rhythm. The American Journal of Psychology, 6(2), 1–46.

Brochard, R., Abecasis, D., Potter, D., Ragot, R., & Drake, C. (2003). The “Ticktock” of Our Internal Clock. Psychological Science, 14(4), 362–366. https://doi.org/10.1111/1467-9280.24441

Buzsáki, G. (2009). Rhythms of the Brain. In Rhythms of the Brain. https://doi.org/10.1093/acprof:oso/9780195301069.001.0001

Cohen, M. X. (2014). Analyzing Neural Time Series Data: Theory and Practice. In MIT Press. https://doi.org/10.1017/CBO9781107415324.004

Colling, L. J., Noble, H. L., & Goswami, U. (2017). Neural entrainment and sensorimotor synchronization to the beat in children with developmental dyslexia: An EEG study. Frontiers in Neuroscience, 11(JUL). https://doi.org/10.3389/fnins.2017.00360

Doelling, K. B., & Poeppel, D. (2015). Cortical entrainment to music and its modulation by expertise. Proceedings of the National Academy of Sciences of the United States of America, 112(45), E6233–E6242. https://doi.org/10.1073/pnas.1508431112

Friston, K. (2005). A theory of cortical responses. Philosophical Transactions of the Royal Society B: Biological Sciences. https://doi.org/10.1098/rstb.2005.1622

Fujioka, T., Trainor, L. J., Large, E. W., & Ross, B. (2012). Internalized Timing of Isochronous Sounds Is Represented in Neuromagnetic Beta Oscillations. Journal of Neuroscience, 32(5), 1791–1802. https://doi.org/10.1523/JNEUROSCI.4107-11.2012

Fujioka, Takako, Ross, B., & Trainor, L. J. (2015). Beta-band oscillations represent auditory beat and its metrical hierarchy in perception and imagery. Journal of Neuroscience. https://doi.org/10.1523/JNEUROSCI.2397-15.2015

Fujioka, Takako, Trainor, L. J., Large, E. W., & Ross, B. (2012). Internalized timing of isochronous sounds is represented in neuromagnetic beta oscillations. Journal of Neuroscience. https://doi.org/10.1523/JNEUROSCI.4107-11.2012

Fujioka, Takako, Zendel, B. R., & Ross, B. (2010). Endogenous Neuromagnetic Activity for Mental Hierarchy of Timing. Journal of Neuroscience, 30(9), 3458–3466. https://doi.org/10.1523/JNEUROSCI.3086-09.2010

Giraud, A. L., & Poeppel, D. (2012). Cortical oscillations and speech processing: Emerging computational principles and operations. Nature Neuroscience, 15(4), 511–517. https://doi.org/10.1038/nn.3063

Goswami, U. (2011). A temporal sampling framework for developmental dyslexia. Trends in Cognitive Sciences, 15(1), 3–10. https://doi.org/10.1016/j.tics.2010.10.001

Grahn, J. A., & Brett, M. (2007). Rhythm and Beat Perception in Motor Areas of the Brain. Journal of Cognitive Neuroscience, 19(5), 893–906. https://doi.org/10.1162/jocn.2007.19.5.893

Grahn, J. A., & McAuley, J. D. (2009). Neural bases of individual differences in beat perception. NeuroImage, 47(4), 1894–1903. https://doi.org/10.1016/J.NEUROIMAGE.2009.04.039

Háden, G. P., Honing, H., Török, M., & Winkler, I. (2015). Detecting the temporal structure of sound sequences in newborn infants. International Journal of Psychophysiology. https://doi.org/10.1016/j.ijpsycho.2015.02.024

Haegens, S., & Zion Golumbic, E. (2018). Rhythmic facilitation of sensory processing: A critical review. Neuroscience and Biobehavioral Reviews, 86(December 2017), 150–165. https://doi.org/10.1016/j.neubiorev.2017.12.002

Jones, M. R. (1976). Time, our lost dimension: Toward a new theory of perception, attention, and memory. Psychological Review, 83(5), 323–355. https://doi.org/10.1037/0033-295X.83.5.323

Jongsma, M. L. A., Desain, P., & Honing, H. (2004). Rhythmic context influences the auditory evoked potentials of musicians and non-musicians. Biological Psychology, 66(2), 129–152. https://doi.org/10.1016/J.BIOPSYCHO.2003.10.002

Kaneshiro, B., Nguyen, D. T., Norcia, A. M., Dmochowski, J. P., & Berger, J. (2020). Natural music evokes correlated EEG responses reflecting temporal structure and beat. NeuroImage, 214, 116559. https://doi.org/10.1016/J.NEUROIMAGE.2020.116559

Koelsch, S., Vuust, P., & Friston, K. (2019). Predictive Processes and the Peculiar Case of Music. In Trends in Cognitive Sciences. https://doi.org/10.1016/j.tics.2018.10.006

Kononowicz, T. W., & van Rijn, H. (2015). Single trial beta oscillations index time estimation. Neuropsychologia, 75, 381–389. https://doi.org/10.1016/J.NEUROPSYCHOLOGIA.2015.06.014

Lakatos, P., Karmos, G., Mehta, A. D., Ulbert, I., & Schroeder, C. E. (2008). Entrainment of neuronal oscillations as a mechanism of attentional selection. Science. https://doi.org/10.1126/science.1154735

Large, E. W., & Jones, M. R. (1999). The dynamics of attending: How people track time-varying events. Psychological Review, 106(1), 119–159. https://doi.org/10.1037//0033-295x.106.1.119

Merchant, H., & Bartolo, R. (2018). Primate beta oscillations and rhythmic behaviors. In Journal of Neural Transmission (Vol. 125, Issue 3, pp. 461–470). Springer-Verlag Wien. https://doi.org/10.1007/s00702-017-1716-9

Merchant, H., Grahn, J., Trainor, L., Rohrmeier, M., & Fitch, W. T. (2015). Finding the beat: A neural perspective across humans and non-human primates. Philosophical Transactions of the Royal Society B: Biological Sciences. https://doi.org/10.1098/rstb.2014.0093

Merker, B. H., Madison, G. S., & Eckerdal, P. (2009). On the role and origin of isochrony in human rhythmic entrainment. Cortex. https://doi.org/10.1016/j.cortex.2008.06.011

Morillon, B., Schroeder, C. E., Wyart, V., & Arnal, L. H. (2016). Temporal prediction in lieu of periodic stimulation. Journal of Neuroscience. https://doi.org/10.1523/JNEUROSCI.0836-15.2016

Mormann, F., Fell, J., Axmacher, N., Weber, B., Lehnertz, K., Elger, C. E., & Fernández, G. (2005). Phase/amplitude reset and theta-gamma interaction in the human medial temporal lobe during a continuous word recognition memory task. Hippocampus, 15(7), 890–900. https://doi.org/10.1002/hipo.20117

Nobre, A.C., Rohenkohl, G., & Stokes, M. (2012). Nervous Anticipation : Top-Down Biasing across Space and Time. Cognitive Neuroscience of Attention.

Nobre, Anna C., & Van Ede, F. (2018). Anticipated moments: Temporal structure in attention. Nature Reviews Neuroscience, 19(1), 34–48. https://doi.org/10.1038/nrn.2017.141

Nozaradan, S. (2014). Exploring how musical rhythm entrains brain activity with electroencephalogram frequency-tagging. In Philosophical Transactions of the Royal Society B: Biological Sciences. https://doi.org/10.1098/rstb.2013.0393

Nozaradan, S., Peretz, I., & Keller, P. E. (2016). Individual differences in rhythmic cortical entrainment correlate with predictive behavior in sensorimotor synchronization. Scientific Reports. https://doi.org/10.1038/srep20612

Nozaradan, S., Peretz, I., Missal, M., & Mouraux, A. (2011). Tagging the Neuronal Entrainment to Beat and Meter. Journal of Neuroscience, 31(28), 10234–10240. https://doi.org/10.1523/JNEUROSCI.0411-11.2011

Nozaradan, S., Schwartze, M., Obermeier, C., & Kotz, S. A. (2017). Specific contributions of basal ganglia and cerebellum to the neural tracking of rhythm. Cortex, 95, 156–168. https://doi.org/10.1016/j.cortex.2017.08.015

Nozaradan, S., Zerouali, Y., Peretz, I., & Mouraux, A. (2015). Capturing with EEG the Neural Entrainment and Coupling Underlying Sensorimotor Synchronization to the Beat. Cerebral Cortex, 25(3), 736–747. https://doi.org/10.1093/cercor/bht261

Obleser, J., Herrmann, B., & Henry, M. J. (2012). Neural oscillations in speech: Don’t be enslaved by the envelope. Frontiers in Human Neuroscience. https://doi.org/10.3389/fnhum.2012.00250

Oostenveld, R., Fries, P., Maris, E., & Schoffelen, J. M. (2011). FieldTrip: Open source software for advanced analysis of MEG, EEG, and invasive electrophysiological data. Computational Intelligence and Neuroscience. https://doi.org/10.1155/2011/156869

Patel, A. D., & Iversen, J. R. (2014). The evolutionary neuroscience of musical beat perception: The Action Simulation for Auditory Prediction (ASAP) hypothesis. Frontiers in Systems Neuroscience. https://doi.org/10.3389/fnsys.2014.00057

Poudrier, È. (2020). The Influence of Rate and Accentuation on Subjective Rhythmization. Music Perception, 38(1), 27–45. https://doi.org/10.1525/mp.2020.38.1.27

Schmidt-Kassow, M., Rothermich, K., Schwartze, M., & Kotz, S. A. (2011). Did you get the beat? Late proficient French-German learners extract strong–weak patterns in tonal but not in linguistic sequences. NeuroImage, 54(1), 568–576. https://doi.org/10.1016/j.neuroimage.2010.07.062

Schroeder, C. E., & Lakatos, P. (2009). Low-frequency neuronal oscillations as instruments of sensory selection. Trends in Neurosciences, 32(1), 9–18. https://doi.org/10.1016/j.tins.2008.09.012

Schröger, E., Kotz, S. A., & SanMiguel, I. (2015). Bridging prediction and attention in current research on perception and action. Brain Research, 1626, 1–13. https://doi.org/10.1016/j.brainres.2015.08.037

Schwartze, M., Keller, P. E., & Kotz, S. A. (2016). Spontaneous, synchronized, and corrective timing behavior in cerebellar lesion patients. Behavioural Brain Research, 312, 285–293. https://doi.org/10.1016/j.bbr.2016.06.040

Schwartze, M., Rothermich, K., Schmidt-Kassow, M., & Kotz, S. A. (2011). Temporal regularity effects on pre-attentive and attentive processing of deviance. Biological Psychology, 87(1), 146–151. https://doi.org/10.1016/j.biopsycho.2011.02.021

Schwartze, M., Stockert, A., & Kotz, S. A. (2015). Striatal contributions to sensory timing: Voxel-based lesion mapping of electrophysiological markers. Cortex, 71, 332–340. https://doi.org/10.1016/j.cortex.2015.07.016

Shalev, N., Nobre, K., & van Ede, F. (2019). Time for What: Breaking Down Temporal Anticipation.

Thut, G., Miniussi, C., & Gross, J. (2012a). The functional importance of rhythmic activity in the brain. In Current Biology (Vol. 22, Issue 16). https://doi.org/10.1016/j.cub.2012.06.061

Thut, G., Miniussi, C., & Gross, J. (2012b). The functional importance of rhythmic activity in the brain. Current Biology, 22(16), R658–R663. https://doi.org/10.1016/j.cub.2012.06.061

Waschke, L., Kloosterman, N. A., Obleser, J., & Garrett, D. D. (2021). Behavior needs neural variability. Neuron, 109(5), 751–766. https://doi.org/10.1016/J.NEURON.2021.01.023

Zoefel, B., ten Oever, S., & Sack, A. T. (2018). The involvement of endogenous neural oscillations in the processing of rhythmic input: More than a regular repetition of evoked neural responses. Frontiers in Neuroscience, 12(MAR), 1–13. https://doi.org/10.3389/fnins.2018.00095

Zoefel, B., & VanRullen, R. (2016). EEG oscillations entrain their phase to high-level features of speech sound. NeuroImage, 124, 16–23. https://doi.org/10.1016/j.neuroimage.2015.08.054

